# Knockout of PI4-Kinase A in GnRH Neurons Causes their Prepubertal Death

**DOI:** 10.64898/2026.02.04.703844

**Authors:** Stephanie Constantin, Naseratun Nessa, Stanko S. Stojilkovic

**Author notes:** Correspondence to Dr. Stephanie Constantin.

## Abstract

The signaling pathways that control embryonic development, migration, and differentiation of gonadotropin-releasing hormone (GnRH) neurons, as well as the postnatal fate, function, and survival of differentiated cells, are the subject of ongoing research. Here, we examined the role of phosphoinositides in this complex multistep process by generating GnRH neuron-specific phosphatidylinositol 4-kinase alpha knockout mice. These mice were healthy and indistinguishable from their control littermates in size. However, adult knockout females and males were infertile due to underdeveloped gonads and reproductive organs. Furthermore, hypothalamic GnRH immunoreactivity was absent, and expression of the hypothalamic *Gnrh1* gene and pituitary gonadotroph-specific genes was reduced. In contrast, hypothalamic kisspeptin immunoreactivity was preserved, and *Kiss1* expression was modified in a nuclei specific-manner, consistent with the loss of circulating sex steroid hormones. Embryonic neurogenesis and migration of GnRH neurons were not impaired, as evidenced by normal *Gnrh1* expression in the hypothalamus of neonatal animals and the presence of immunoreactive GnRH neurons in infantile mice in comparable number and distribution to age-matched controls. However, their cellular degeneration was evident, accompanied by reduced *Gnrh1* expression. GnRH neuron-specific tdTomato expression confirmed their postnatal degeneration and death, whereas ectopic tdTomato cells located in the lateral septum remained unaffected. Together, these findings indicate that phosphoinositides dependent on phosphatidylinositol 4-kinase alpha activity are not critical for embryonic steps in the development of the GnRH neuronal network, but are essential for the postnatal function and survival of these cells.

**Significance Statement:** Differentiation of neuroendocrine GnRH cells involves neurogenesis in the olfactory placodes, migration to the hypothalamus, projection to the median eminence, and connections with upstream neurons, including kisspeptin neurons. Here we show that knockout of phosphatidylinositol 4-kinase alpha in GnRH neurons does not affect these strps of embryonic development. However, the activity of this enzyme is essential for postnatal survival of GnRH neurons; in the absence of this gene, the neurons die, causing infertility in both female and male mice.

## Introduction

Gonadotropin-releasing hormone (GnRH)-secreting neurons originate in the medial olfactory placodes and migrate to the hypothalamus during embryonic development (Duittoz et al., 2022). Their cell bodies form a rostrocaudal continuum from the preoptic area to the mediobasal hypothalamus, with the highest cell density around the organum vasculosum lamina terminalis (OVLT) and projecting to the median eminence. During in utero development, connections between GnRH neurons and kisspeptin neurons are establish in the arcuate nucleus (ARN) (Kumar et al., 2014). Before the onset of puberty, connections between these two cell types are also established in the rostral periventricular area of the third ventricle (RP3V) (Semaan et al., 2013), as the responsiveness of GnRH neurons to kisspeptin increases (Han et al., 2005). Kisspeptin neurons in the ARN, also known as KNDy neurons, coexpress neurokinin B and dynorphin A (Lehman et al., 2010). These neurons act as GnRH pulse generator by innervating distal projections of GnRH neurons in the median eminence in both females and males (Wang et al., 2020), and their function is inhibited by gonadal steroids. RP3V kisspeptin neurons project to GnRH cell bodies in the preoptic area and to the distal projections of GnRH neurons (Wang et al., 2020) and act as GnRH surge generator. Unlike KNDy neurons, RP3V kisspeptin neuron function is stimulated by gonadal steroids, leading to preovulatory GnRH release (Clarkson et al., 2008). Pulsatile release of GnRH stimulates expression of the pituitary gonadotropin genes *Fshb* and *Lhb*, their protein synthesis, and the release of follicle-stimulating hormone and luteinizing hormone. These hormones then control gametogenesis and steroidogenesis in the gonads of both sexes and ovulation in females (Constantin et al., 2022).

The signaling pathway involved in communication between GnRH neurons and kisspeptin neurons is well characterized. Kisspeptin receptors expressed in GnRH neurons signal via the Gq/phospholipase C pathway, which leads to the hydrolysis of phosphatidylinositol 4,5-bisphosphate (PI(4,5)P2) to inositol 1,4,5-trisphosphate and diacylglycerol (Liu et al., 2008). This is followed by the initiation of inositol 1,4,5-trisphosphate receptor-dependent release of calcium from intracellular pools and modulation of potassium and nonselective cation channels (Zhang and Spergel, 2012). This causes depolarization of GnRH neurons accompanied by an increase in firing frequency (Constantin et al., 2013) and/or facilitation of voltage-gated calcium influx (Iremonger et al., 2017). A decrease in PI(4,5)P2 levels during this signaling cascade prevents the subsequent kisspeptin response, but this refractory period can be reduced by replenishing PI(4,5)P2 (Constantin et al., 2021). Therefore, PI(4,5)P2 is a key player and regulator of the kisspeptin-induced response in GnRH neurons. However, the roles of this and other phosphoinositides in the development of the GnRH neuronal system and its function have not been studied.

Phosphoinositides are formed by the regulated activity of phosphoinositide kinases and phosphatases. Among these enzymes, phosphatidylinositol 4-kinase alpha (PI4KA) is responsible for the formation of phosphatidylinositol 4-phosphate (PI4P), which is itself a signaling molecule and the starting point for the formation of PI(4,5)P2 in the plasma membrane. PI(4,5)P2 is vital for numerous cellular functions, including phospholipase C signaling, ion channel gating, and control of exocytotic pathways. Furhtermore, phosphorylation of PI(4,5)P2 by class I phosphatidylinositol-3-kinase generates phosphatidylinositol 3,4,5-trisphosphate (PI(3,4,5)P3), which exerts its intracellular messenger functions by binding to numerous effectors containing a pleckstrin homology domain (Stojilkovic and Balla, 2023). PI4KA is also well expressed in GnRH neurons (Burger et al., 2018), but its specific function has not been studied in these cells. Global knockout of PI4KA in mouse models is embryonically lethal (Nakatsu et al., 2012), but cell type–specific PI4KA knockout experiments offer the potential to study the effects of this gene on specific cells function (Constantin et al., 2023). Here, we present data obtained from experiments with GnRH neuron-specific knockout mice to determine the potential role of PI4KA during embryonic development of these neurons, as well as on their postnatal fate and function.

## Material and Methods

### Animals

All experimental procedures were approved by the National Institute of Child Health and Human Development, Animal Care and Use Committee (Protocol 22-041). Pi4ka-LoxP mice (MGI:6294260) (Bojjireddy et al., 2014) were crossed with Rosa-tdTomato mice (MGI:3813512) (Madisen et al., 2010) to generate dams Pi4ka^LoxP/LoxP^ Rosa^tdTom/tdTom^. Pi4ka-LoxP mice were crossed with Gnrh1-cre mice (MGI: 3691288) (Yoon et al., 2005) to generate Gnrh1^Cre/Cre^ Pi4ka^LoxP/wt^ bucks. The crossing resulted in conditional knockout mice in which PI4KA was specifically inactivated in GnRH neurons (Gnrh1^Cre/wt^ Pi4ka^LoxP/LoxP^ Rosa^tdTom/wt^; hereafter knockouts) and their control littermates (Gnrh1^Cre/wt^ Pi4ka^LoxP/wt^ Rosa^tdTom/wt^; hereafter controls). Cre-dependent recombination could be monitored with tdTomato expression in both knockouts and controls. Fertility was assessed as follows: three to five knockout and control mice (7–12-week-old) were mated with wild-type C57/Bl6 of the opposite sex for <4 weeks to assess their fertility. The time of the first litter, relative to the date of the mating, was recorded. Necropsy was performed at postnatal day (PND) 45 on control and knockout mice.

### qRT-PCR analysis

Neonatal (PND3), infantile (PND 9 and PND10), juvenile (PND15 and PND20), peripubertal (PND45), and adult (8 to 25 weeks) females and males were used to collect hypothalamic and pituitary tissues. Before PND10, the whole hypothalamus was collected. From PND10, the hypothalamus was divided into rostral and caudal parts using the optic tract as a demarcation. Whole pituitaries were collected simultaneously. All tissues were snap frozen on dry ice. Hypothalamic and pituitary RNA were isolated using the RNeasy Plus Mini Kit (Qiagen, Valencia, CA), and reverse transcription was performed using the Transcriptor First Strand cDNA Synthesis Kit (Roche Applied Sciences, Indianapolis, IN), according to the manufacturer’s respective protocols. Applied Biosystems predesigned TaqMan Gene Expression Assays (Applied Biosystems, Waltham, MA) were used for the following genes: *Gapdh* (Mm99999915_g1), *Gh1* (Mm00433590_g1), *Gnrh1* (Mm01315604_m1), *Gnrhr* (Mm00439143_m1), *Kiss1* (Mm03058560_m1), *Lhb* (Mm01205505_g1), *Mkrn3* (Mm00844003_s1), *Pdyn* (Mm00457573_m1), *Pomc* (Mm00435874_m1), *Prl* (Mm00599950_m1), *Spp1* (Mm00436767_m1), *Tac2* (Mm01160362_m1), and *Tshb* (Mm03990915_g1).

### Immunolabeling

Mice, anesthetized with a ketamine/xylazine cocktail (200/20 mg/kg), were transcardially perfused with 0.1 M phosphate buffer saline (PBS), followed by PBS containing 4% formaldehyde. Brains were postfixed overnight and then transferred to 30% PBS containing sucrose for 2 nights. Brains were snap-frozen on dry ice using Tissue-Plus OCT (Fisher Healthcare/ThermoFisher Scientific) and stored at ™80 °C until sectioning. Adult brains were cut into four sets of 40 µm coronal sections using a Leica SM2010 R sliding microtome (Wetzlar, Germany). PND10 brains were cut into two sets of 50 µm coronal sections. All sections were stored at -20 °C in cryoprotectant until staining (>5 days). For immunochemistry, PND10 and adult free-floating sections were processed for GnRH and kisspeptin with rabbit anti-GnRH antibody (RRID: AB_572248; dilution 1:15,000) or kisspeptin antibody (RRID:AB_2314709; dilution 1:10,000), as previously described (Sokanovic et al., 2023). For immunofluorescence, PND10 and adult free-floating sections were incubated for 1 h at room temperature in blocking solution (10% normal goat serum plus 0.3% Triton X-100), washed several times in PBS, and incubated (2–3 nights, 4 °C) in guinea pig anti-GnRH antibody (RRID: AB_2721118). Sections were then washed in PBS, incubated (1 h, room temperature) with a secondary AlexaFluor 488 goat anti-guinea pig antibody (AB_2534117; 1:1,000 in PBS/0.3% Triton X-100). All images were captured using 10X and 40X magnification objectives on an Eclipse Ti2 microscope (Nikon, Melville, NY). Excitation was provided by a SOLA V-nIR light system (Lumencor, Beaverton, OR), and emission was captured by an ORCA-Flash4.0 V3 digital camera (Hamamatsu, Bridgewater, NJ). Images were processed using FIJI (Schindelin et al., 2012) and montages were created in Photoshop (Adobe, San Jose, CA).

### Statistical analysis

Results are presented as representative histological images or mean ± SEM values from at least three similar experiments, and the number of replicates is indicated in Results or Figure Legends. Graphs were generated and statistically analyzed using the KaleidaGraph program (Synergy Software, Reading, PA) with one-way ANOVA and post hoc multiple comparison test for data containing more than two groups, with at least P < 0.01 considered statistically significant. Figures were finalized using Adobe Photoshop and Illustrator (Adobe, San Jose, CA).

## Results

### GnRH neuron-specific PI4KA knockout mice are infertile

The experiments were conducted with female and male wild-type, control, and knockout mice. Control and knockout mice were healthy, and their developmental pattern was indistinguishable from wild-type mice. After housing with wild-type mice of the opposite sex, control females had their first litter within 26 ± 3 days (5 breeding pairs) and control males had their first litter within 23 ± 2 days (7 breeding pairs). However, neither knockout females nor males had litters when mated with wild-type mice of the opposite sex for >4 weeks.

The infertility of the knockouts was associated with the status of their reproductive organs, as shown in Fig. 1*A1, A2, B1*, and *B1* for females and Fig. 1 *A3, A4, B3*, and *B1* for males; controls (top panels) and knockouts (bottom panels), all PND45. In contrast to controls, knockouts lacked external features typically associated with sexual maturity; knockout females did not show a vaginal opening (Fig. 1*A1* vs *B1*) and males did not show separation of the prepuce, also known as foreskin separation (Fig. 1*A3* vs *B1*). Necropsy further revealed that knockout females (Fig. 1*B1* vs *B1*) and males (Fig. 1*B1* vs *B1*) displayed underdeveloped reproductive organs compared with their control littermates. These data clearly indicate that the hypothalamic-pituitary-gonadal axis is not operational in PI4KA knockouts.

**Figure 1.**
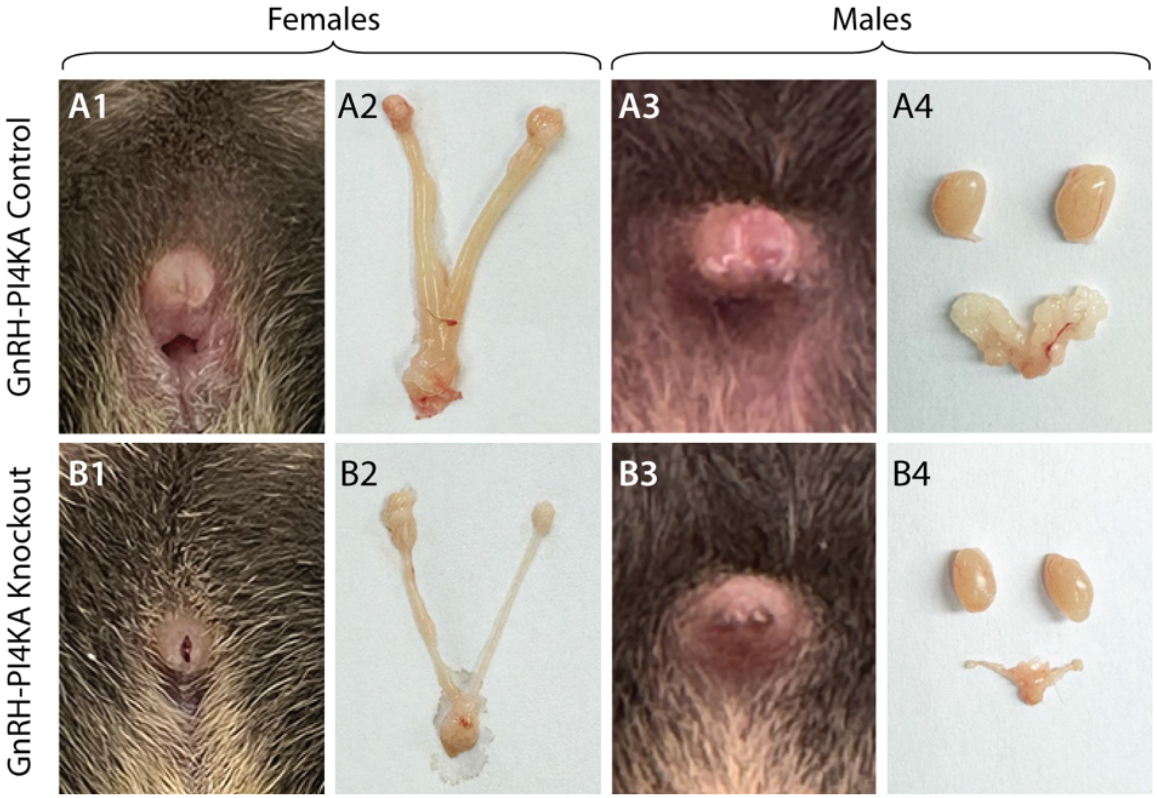
Comparison of reproduc0ve organs of control and knockout mice at seven weeks of age. In control females, the vagina was perforated (***A1***) and the animals had a normally developed uterus and ovaries (***A2***). Control males showed foreskin separa0on (***A3***), and the testes and seminal vesicles were normally developed (***A4***). In contrast, female knockouts showed imperforate vaginas (***B1***) and underdeveloped uterus and ovaries (***B2***). Male knockouts did not show foreskin separa0on (***B3***), and the testes and seminal vesicles were atrophied (***B4***).

### Loss of GnRH expression causes infertility of knockout mice

Because PI4KA knockout is specific for GnRH-secreting cells, and hypothalamic GnRH-secreting neurons are essential for the establishment and function of the hypothalamic-pituitary-gonadal axis, in further experiments we performed immunostaining to assess GnRH peptide expression in hypothalamic tissue of adult control and knockout mice as described in Material and Methods. Immunostaining was examined across the rostrocaudal GnRH continuum. For representation, we focused on GnRH immunoreactivity around the OVLT area (Fig. 2*A-D*), which contains many GnRH neurons. Adult control females (Fig. 2*A*) and males (Fig. 2*C*) showed a typical distribution of GnRH-expressing cell bodies. In contrast, knockout females (Fig. 2*B*) and males (Fig. 2*D*) had no GnRH immunoreactivity.

**Figure 2.**
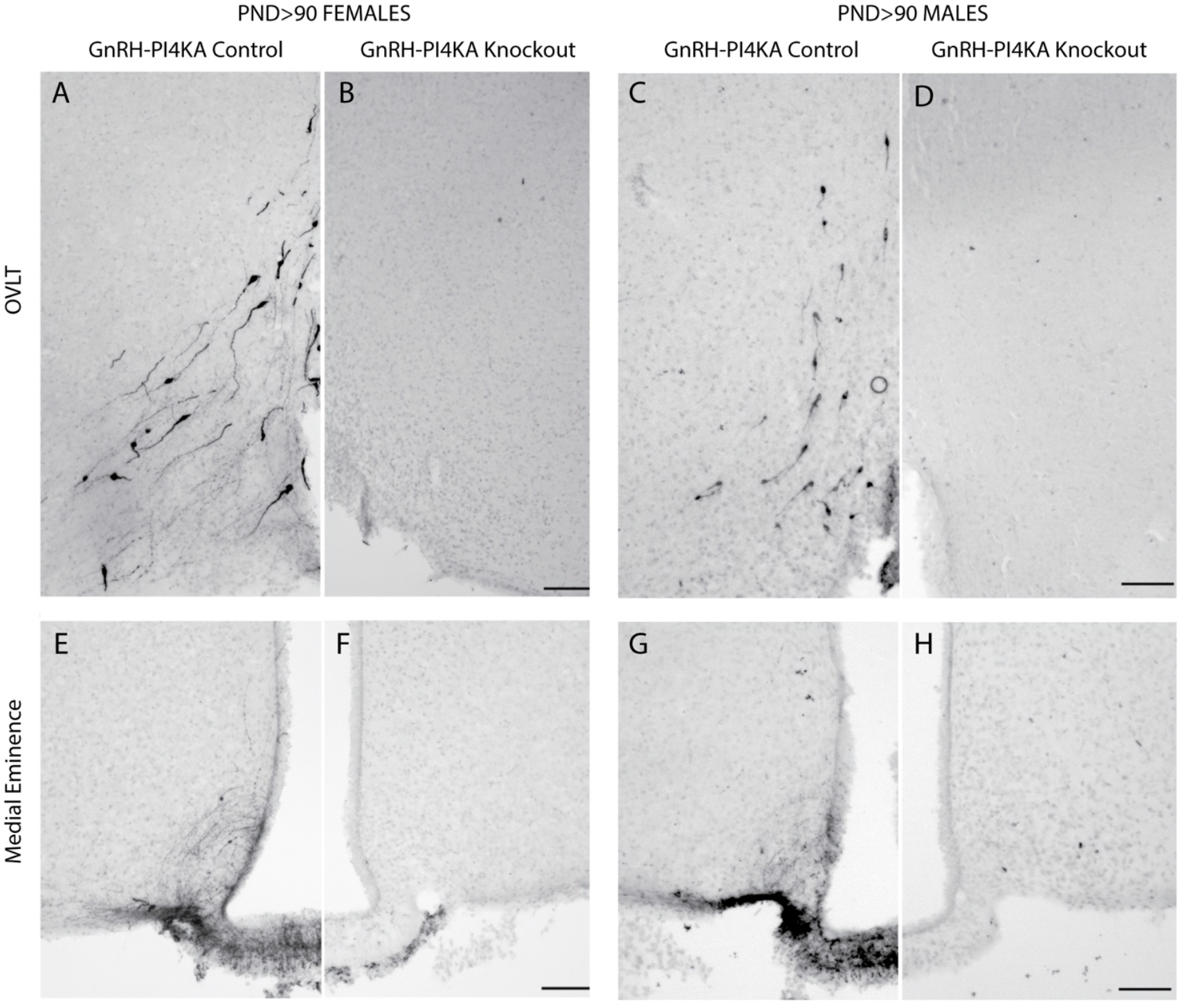
Immunostaining of GnRH-expressing neurons in hypothalamic 0ssue of adult control and knockout mice. ***A*** to ***D***, GnRH-expression paDern in the organum vasculosum lamina terminalis (OVLT) of control (***A***) and knockout (***B***) females, and control (***C***) and knockout (***D***) males. Typical bipolar GnRH-posi0ve neurons were present in control females and males, but were not visible in the OVLT area of knockouts. ***E*** to ***H***, GnRH expression paDern in the medial eminence of control (***E***) and knockout (***F***) females, and control (***G***) and knockout (***H***) males. Typical GnRH-posi0ve fibers were detected in the medial eminence of controls, but not of knockouts. Horizontal bars at 100 µM.

We also analyzed GnRH immunoreactivity in median eminence of adult control and knockout animals (Fig. 2*E-H*). In general, GnRH neurons from the OVLT project long processes to the median eminence at the base of the hypothalamus, where they branch extensively and form terminals near blood vessels to release the hormone into the pituitary portal system (Herde et al., 2013). GnRH immunoreactivity highlighted fibers in control females and males (Figs. 2*E* and *B2*), but not in knockout females and males (Fig. 2*F* and *B2*).

By binding to its receptor in pituitary gonadotrophs, GnRH controls the expression of gonadotroph-specific *Lhb, Fshb*, and *Gnrhr* (Constantin et al., 2022). Therefore, the expression of these genes in the pituitary can be used as an indicator of the status of GnRH secretion. Consistent with the lack of hypothalamic immunoreactivity for GnRH, the expression of *Lhb* and *Gnrhr* was significantly reduced in knockout females and males compared with adult controls. The expression of *Spp1*, another gonadotroph-specific gene within the pituitary cell populations (Fletcher et al., 2019), was also significantly reduced in knockout females and males (Fig. 3*A*). Expression of *Pomc, Tshb*, and *Gh1*, the marker genes of corticotroph/melanotrophs, thyrotrophs and somatotrophs, respectively, was unaffected, while expression of *Prl*, a marker gene of lactotrophs, was significantly reduced in knockout females and males (Fig. 3*B*). Together, the lack of immunoreactive GnRH neurons and the reduced expression of *Lhb* and *Gnrhr* in knockouts provide a rationale for the underdeveloped reproductive organs and infertility of these animals, and the concomitant reduction in gonadal steroidogenesis leads to a decrease in *Spp1* and *Prl* expression.

**Figure 3.**
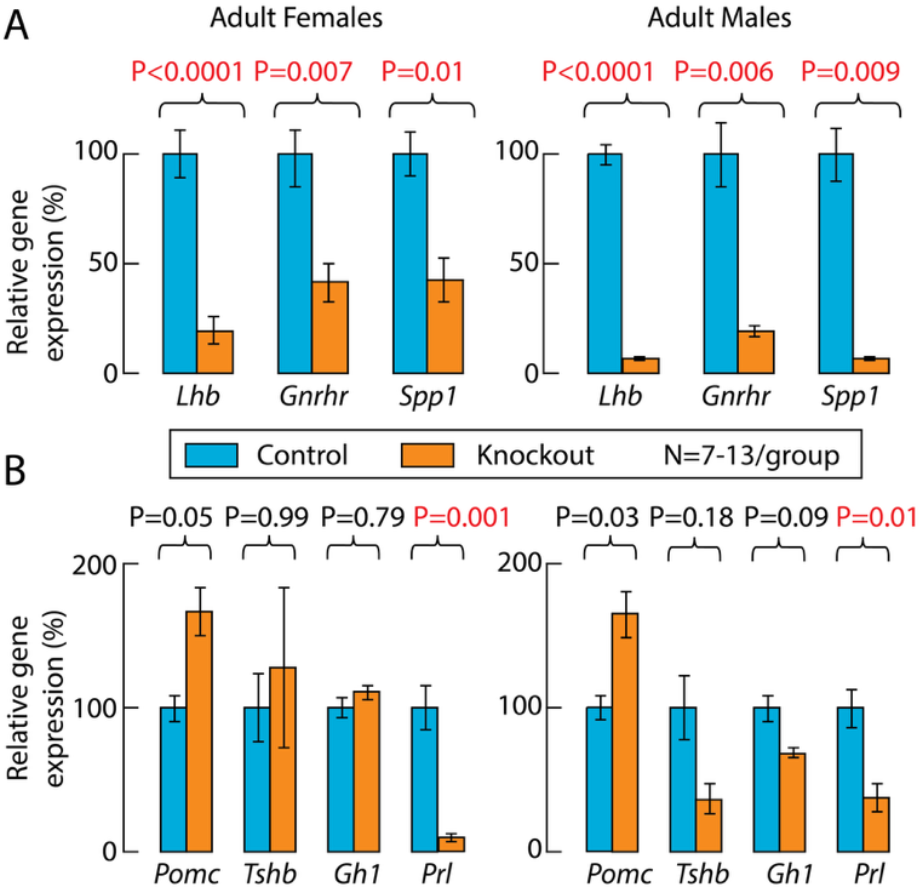
Expression of pituitary genes in adult control and knockout mice. ***A***, Expression of gonadotroph-specific genes *Lhb, Gnrhr*, and *Spp1*. ***B***, Expression of cor.cotroph/melanotroph-specific *Pomc*, thyrotroph-specific *Tshb*, somatotroph-specific *Gh1*, and lactotroph-specific *Prl*. Gene expression was assessed by qRT-PCR using whole pituitary for mRNA extrac0on, values for controls are shown as 100%, and for knockouts as percentage change rela0ve to controls. Data are presented as means ± SEM, sta0s0cal analysis was performed using data before normaliza0on, and P values were calculated using Welch’s ANOVA followed by post hoc DunneD’s T3 mul0ple comparisons. Red numbers indicate significant di’erences.

### Hypothalamic kisspeptin immunoreactivity is preserved in adult knockout mice

It is well known that kisspeptin neurons provide a key stimulatory input to GnRH neurons (Han et al., 2015), and that *Kiss1* expression is stimulated or inhibited by circulating sex steroid hormones, depending on the specific brain region (Smith et al., 2005a; Smith et al., 2005b). Due to the lack of detectable GnRH immunoreactivity in knockouts, which is associated with underdeveloped gonads, we also analyzed the status of kisspeptin neurons. Figure 4 shows that both controls and knockouts express two kisspeptinergic subpopulations: one located in RP3V (A - D) and another in the ARN (E - H). RP3V kisspeptin expression was sexually dimorphic; it was highly expressed in control females (Fig. 4A and B) and weakly expressed in males (Fig. 4C and D). The density of fibers was reduced in knockout females, but was less pronounced in males due to minimal RP3V kisspeptin immunoreactivity. In contrast, ARN kisspeptin expression was not sexually dimorphic but was affected by PI4KA knockout in GnRH neurons of females (Fig. 4E vs F) and males (Fig. 4G vs H). In both sexes, controls showed fiber-rich immunoreactivity in ARN without identifiable kisspeptin cell bodies, whereas knockouts showed fiber-poor immunoreactivity with readily identifiable kisspeptin cell bodies. Thus, the loss of GnRH in knockouts does not directly reflect changes in the population of kisspeptin neurons, but rather the loss of GnRH-dependent gonadal function causes changes in the pattern of kisspeptin expression.

**Figure 4.**
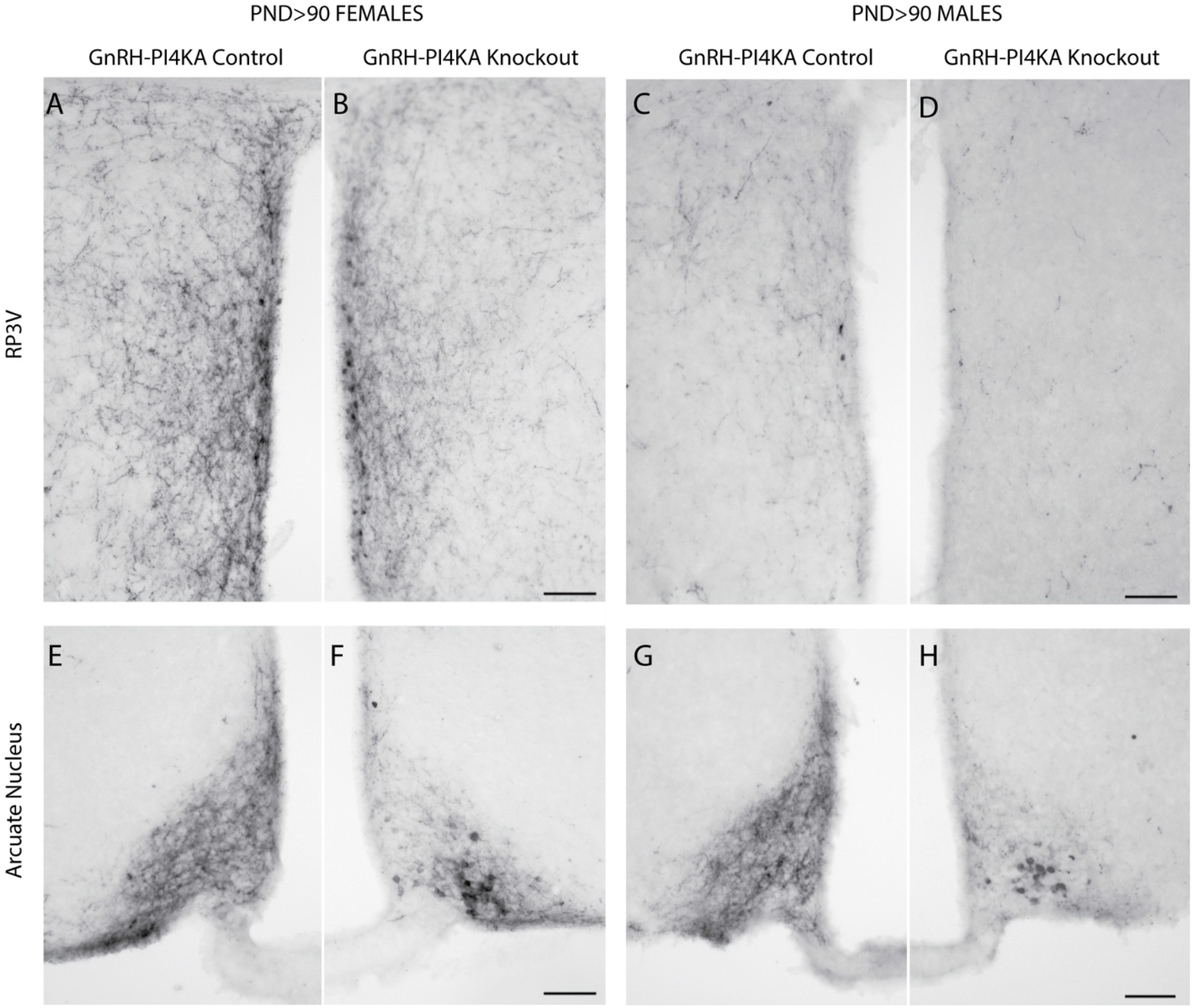
Kisspep0n immunoreac0vity in hypothalamic 0ssue of adult control and knockout mice. ***A*** to ***D***, The expression paDern of kisspep0n in the rostral periventricular area of the third ventricle (RP3V) of control (***A***) and knockout (***B***) females, and of control (***C***) and knockout (***D***) males. ***E*** to ***H***, The expression paDern of kisspep0n in the arcuate nucleus of control (***E***) and knockout (***F***) females, and of control (***G***) and knockout (***H***) males. Horizontal bars at 100 µM.

### *Gnrh1* expression is lost postnatally in knockout mice

The differentiation of GnRH neurons in the olfactory placodes and their migration to the hypothalamus are restricted to embryonic life (Duittoz et al., 2022). Therefore, the loss of GnRH peptide expression in adult knockout mice may reflect a loss of neurogenesis, or neuronal migration embryonically, or changes in neuronal structure and function postnatally. To address this issue, we studied *Gnrh1* expression by qRT-PCR using whole hypothalamic tissue from neonatal animals (PND3) and infantile animals (PND9). At PND3, *Gnrh1* expression was comparable in control and knockout females, but was significantly reduced in knockout females at PND9, while *Kiss1* expression remained unchanged at both ages (Fig. 5A). The same conclusion was reached in experiments with PND3 and PND9 control and knockout males (Fig. 5*B*).

**Figure 5.**
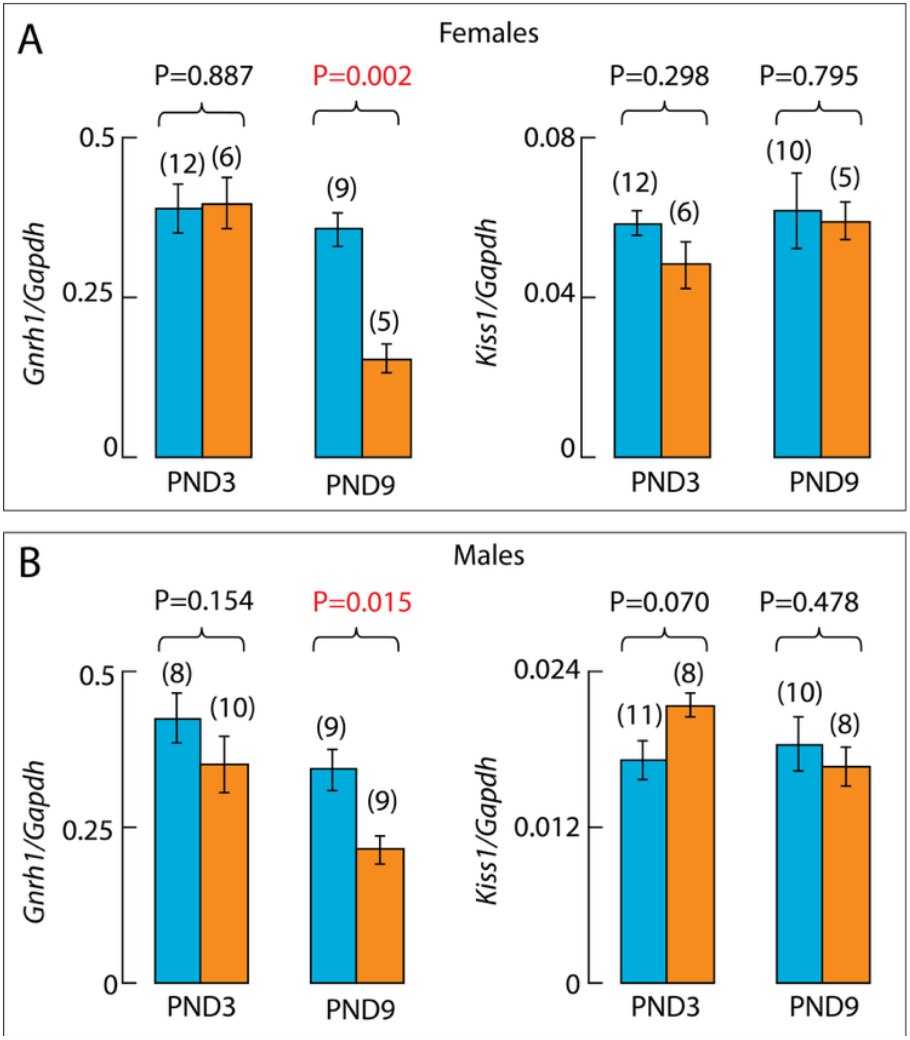
qRT-PCR analysis of Gnrh1 and Kiss1 gene expression in the hypothalamus of control (blue) and knockout (orange) mice. ***A*** and ***B***, *Gnrh1* (***le0***) and *Kiss1* (***right***) expression assessed in the whole hypothalamus of neonatal (PND3) and infan0le (PND9) controls and knockout females (***A***) and males (***B***). Data are presented as means ± SEM and numbers in parentheses indicate the number of animals per group. P values were calculated using one-way ANOVA followed by post hoc Fisher’s least significant di?erence as described in Methods. Red numbers indicate significant di?erences.

In further experiments, hypothalamic GnRH immunoreactivity was assessed at PND10 when *Gnrh1* expression was significantly reduced but not abolished in knockouts compared with age-matched controls. As in adult controls (Fig. 2), this analysis was illustrated with GnRH immunoreactivity around the OVLT (Fig. 6*A-D*), and in the median eminence (Fig. 6*E-H*). Control females (Fig. 6*A*) and males (Fig. 6*C*) showed a distribution of unipolar and bipolar neurons in the hypothalamus comparable to adult controls (Fig. 2*A* and *B2*). PND10 knockouts displayed a similar distribution of GnRH-immunoreactive cells (Fig. 6*B* and *B6*), in contrast to adult knockouts (Fig. 2*B* and *B2*). However, in PND10 knockouts, most GnRH cell bodies appeared round without processes. Furthermore, GnRH immunoreactivity was present but somewhat reduced in the median eminence of knockouts, compared with age-matched controls (Fig. 6*E-H*). Together, this suggests that differentiation and migration of GnRH neurons are not impaired in knockouts, but *Gnrh1* expression is lost postnatally.

**Figure 6.**
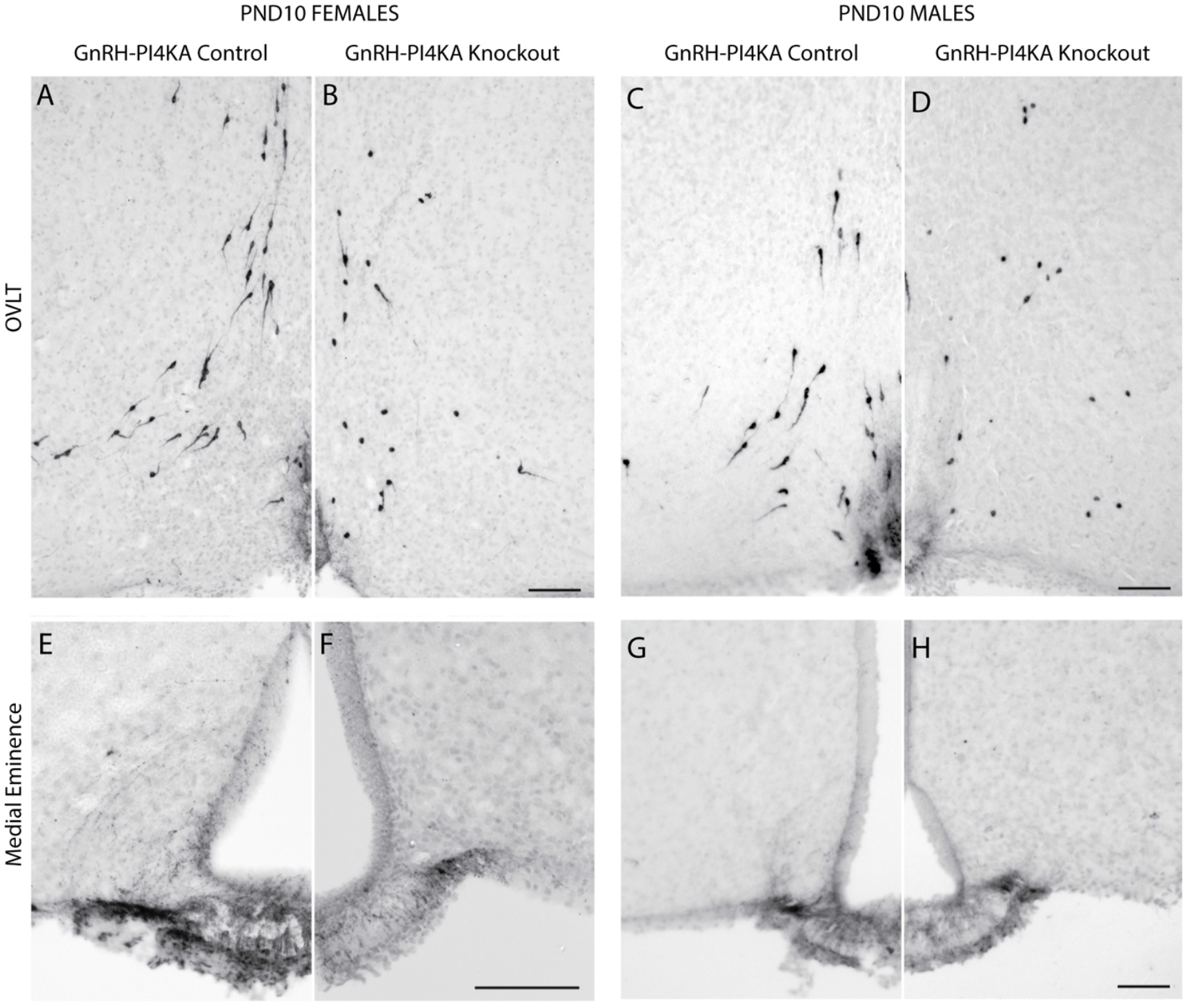
Immunostaining of GnRH neurons in hypothalamic 0ssue from infan0le control and knockout mice. ***A*** to ***D***, GnRH-posi0ve neurons in the OVLT hypothalamic area of control (***A***) and knockout (***B***) females, and control (***C***) and knockout (***D***) males. Note the typical distribu0on of bipolar GnRH-posi0ve neurons in control females and males. GnRH-posi0ve neurons in knockouts showed a similar distribu0on paDern, but cells were predominantly round. ***E*** to ***H***, GnRH expression paDern in the medial eminence of control (***E***) and knockout (***F***) females, and control (***G***) and knockout (***H***) males. It should be noted that GnRH immunoreac0vity was reduced in medial eminence of knockouts. Horizontal bars at 100 µM.

### Opposing effects of knockout on *Gnrh1* and *Kiss1* expression during development

In further experiments, hypothalamic tissue was divided into rostral and caudal parts to examine the kinetics of postnatal *Gnrh1* and *Kiss1* expression in the two regions. Experiments were conducted with females and males aged 10, 15, 20, 45 and 90 days postpartum. Data from females and males were separated to monitor the status of kisspeptinergic neuronal subpopulations. The methodology was evaluated by comparing *Gnrh1* expression in the rostral (including OVLT+RP3V areas) and caudal (including ARN) regions. As expected, using Welch’s ANOVA followed by post hoc Dunnett’s T3 multiple comparisons test, *Gnrh1* expression was significantly higher in the rostral hypothalamus than in the caudal hypothalamus in females (1.090±0.136 vs 0.196±0.057; P = 0.0005) and males (0.798±0.074 vs 0.150±0.058; P = 0.0034). *Kiss1* expression was significantly higher in the rostral hypothalamus in females than males (0.355±0.054 vs 0.039±0.004; P = 0.0014), while *Kiss1* expression was similar in the caudal hypothalamus in females and males (0.106±0.027 vs 0.046±0.010; P = 0.2157).

Figure 7*A* and *B7* show the developmental profile of *Gnrh1* expression in the rostral hypothalamus. In the control group, *Gnrh1* expression was at comparable levels in females and males, but with somewhat different developmental profiles. In females, there was a prepubertal increase in expression of this gene, which was not evident in males. In the knockouts, there was a progressive loss of *Gnrh1* expression in both females and males, which was virtually undetectable in PND45 and PND90. In all ages, a difference between *Gnrh1* expression in the controls and knockout groups was highly significant (P< 0.0001). These results clearly indicate that the loss of GnRH immunoreactivity in adult knockouts reflects a reduction in *Gnrh1* expression in both sexes.

**Figure 7.**
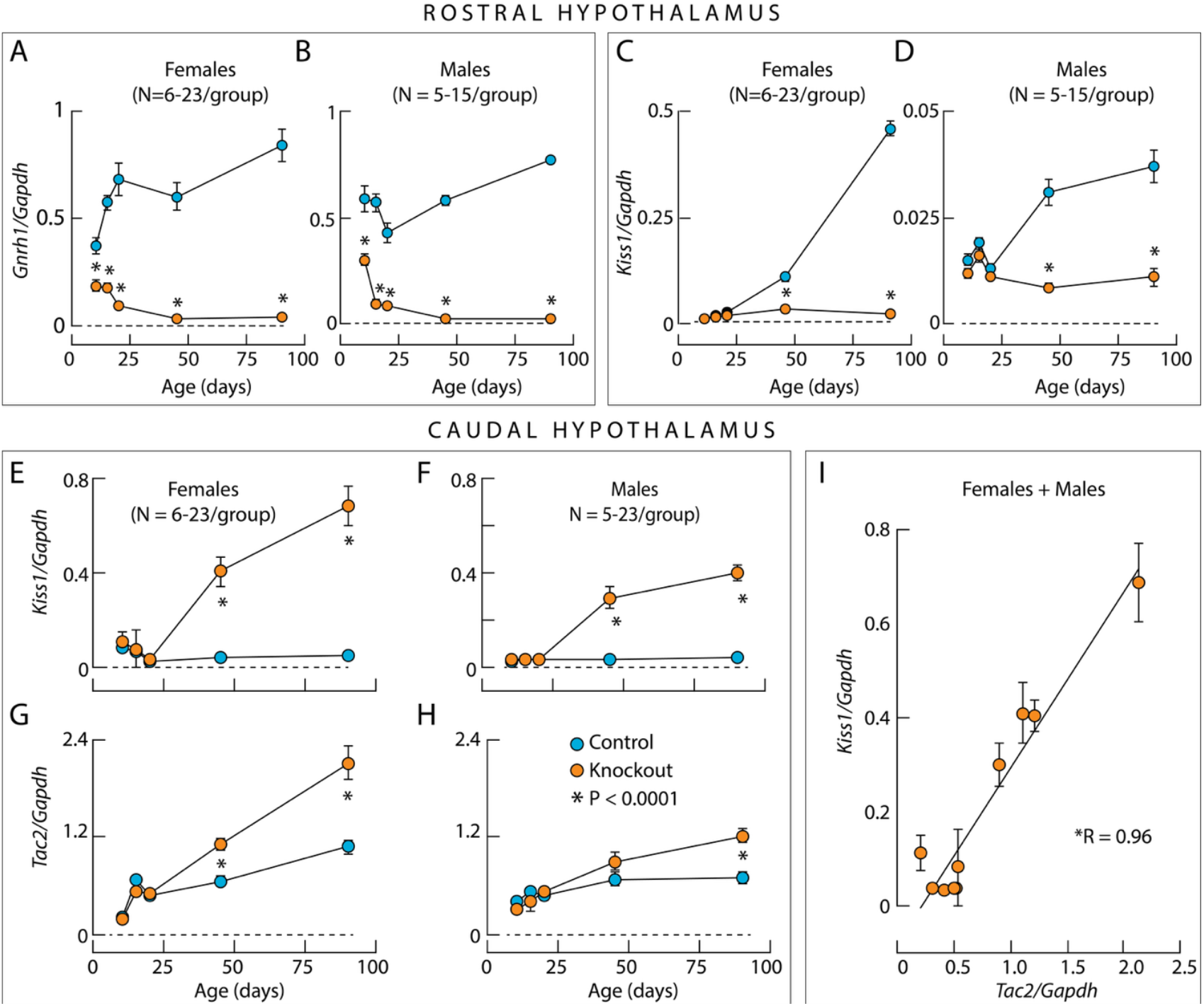
Developmental gene expression profile in the rostral and caudal hypothalamus of control and knockout mice. ***A*** to ***D***, *Gnrh1* expression (***A*** and ***B***) and *Kiss1* expression (***C*** and ***D***) in the rostral hypothalamus of both sexes. Note the 10-fold di?erence in *Kiss1 expression of* control and knockout female (***C***) and male (***D***) mice. ***E*** to ***H***, Facilita0on of *Kiss1* expression (***E*** and ***F***) and *Tac2* expression (***G*** and ***H***) in female (***E*** and ***G***) and male (***F*** and ***H***) knockout mice in the caudal hypothalamus. ***I***, Linear correla0on between *Tac2* and *Kiss1* expression in the caudal hypothalamus of female and male knockout mice. Asterisks indicate significant (P < 0.0001) di?erences between pairs (***A*** to ***H***) using post hoc Šídák’s mul0ple comparisons test ader one-way ANOVA, and for the correla0on coe?cient R using linear regression calcula0on. The legend in ***H*** applies to all panels.

Unlike *Gnrh1* expression, *Kiss1* expression in the rostral hypothalamus was sex-specific, with approximately 10-fold higher expression in females than in males. In both sexes, there was a significant increase in *Kiss1* expression with age (Fig. 7C and *B7*). *Kiss1* expression was similar in knockout mice and control mice up to PND20, but *Kiss1* expression was not increased in knockout females at PND45 and PND90. In the caudal hypothalamus of both sexes, *Kiss1* expression was similar in control and knockout mice up to PND20. However, in knockouts, *Kiss1* expression increased significantly above control levels at PND45 and PND90 (Fig. 7*E* and *B7*).

Since ARN kisspeptin neurons coexpress *Tac2* and *Pdyn* to drive their autonomous activity to generate GnRH pulses (Han et al., 2023), the expression of these genes was also examined. In both sexes, *Tac2* expression was similar up to PND20 in knockout and control groups. At PND45, *Tac2* expression was increased in knockout females compared to control females, but not in knockout males (Fig. 7*G* and *B7*), whose *Kiss1/Tac2* neuronal development was delayed compared to females (Kauffman et al., 2009). Accordingly, at PND90, there was a significant increase in *Tac2* expression in males as well. Moreover, there was a highly significant correlation between *Tac2* and *Kiss1* expression (Fig. 7*I*). In both sexes, *Pdyn* expression was similar between controls and knockouts at all ages studied, indicating that its expression was not affected by gonadal steroids (data not shown). These experiments suggest that the expression pattern of *Kiss1* and *Tac2*, but not *Pdyn* expression, reflects the status of gonadal steroidogenesis. Notably, in both sexes, the expression of *Mkrn3* in hypothalamic tissue decreased during postnatal maturation with similar kinetics between controls and knockout groups (data not shown). This indicates that *Mkrn3* expression, which is crucial for regulating the onset of puberty (Abreu et al., 2013), is independent of gonadal steroid hormones and *Gnrh1*, the latter being consistent with the literature (Abreu et al., 2020).

### Hypothalamic GnRH neurons have altered morphology in infantile knockout mice

Because GnRH immunoreactivity is associated with *Gnrh1* expression, the differences in shape of PI4KA knockout GnRH neurons may reflect a reduction in GnRH immunoreactivity in processes below detection by immunostaining rather than actual changes in the morphology of these cells. To test this hypothesis, we visualized GnRH neurons using the cell type-specific expression of tdTomato, which is known to be ubiquitously expressed after Cre recombination of the Rosa locus and to fill the entire cell. Experiments were performed in control and knockout females. In control group, cells were unipolar or bipolar with similar shapes when immunolabeled with GnRH antibody (Fig. 8*A*), highlighted with tdTomato (Fig. 8*B*), or when merged (Fig. 8*C*). In the knockout group, both GnRH antibody (Fig. 8*D*), and tdTomato (Fig. 8*E*) highlighted the round cell bodies of knockout neurons, confirming changes in cell morphology during the postnatal period. Therefore, it is reasonable to conclude that PI4KA knockout in hypothalamic GnRH neurons leads to their postnatal regression.

**Figure 8.**
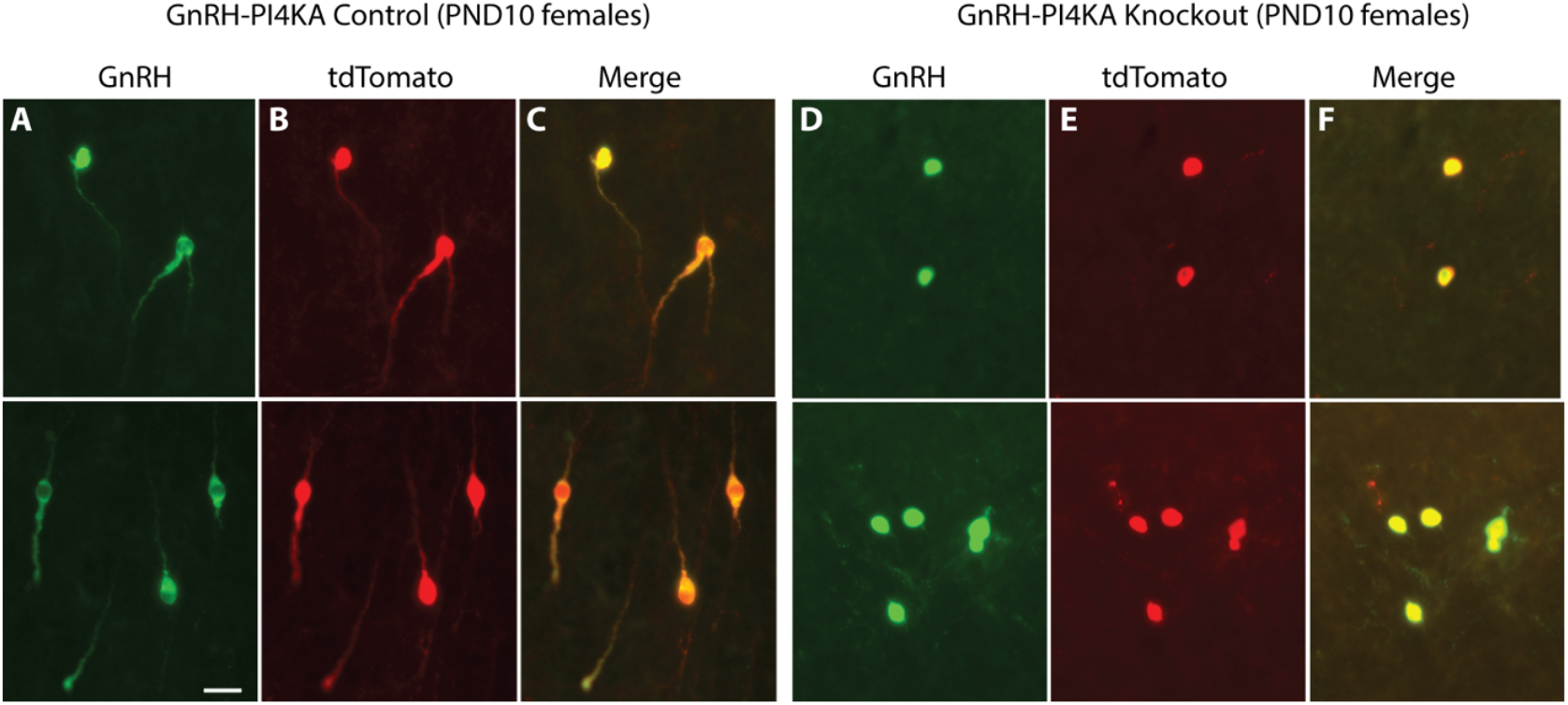
Dual iden0fica0on of GnRH neurons in hypothalamic 0ssue from infan0le control and knockout female mice. GnRH neurons were iden0fied by GnRH immuno?uorescence (green) and cell-type-specific tdTomato expression. For details, see Material and Methods. Representa0ve images from the OVLT area are shown (upper and lower panels). ***A*** to ***C***, Expression of GnRH (***A***), tdTomato (***B***) and their merge (***C***) in control mice. ***D*** to ***F***, Expression of GnRH (***D***), tdTomato (***E***) and their merge (***F***) in knockout mice. Note the same shape of cells visualized by GnRH and tdTomato ?uorescence in control and knockout mice. The horizontal bar at 20 μm applies to all panels.

### GnRH neurons, but not ectopic tdTomato cells, are lost in adult knockouts

During development, some cells, not derived from the olfactory placodes, transiently express *Gnrh1* in the lateral septum, and other areas (Skynner et al., 1999). This transient expression is sufficient to induce *Gnrh1*-driven Cre expression, *Rosa* recombination, and tdTomato expression in cells other than neuroendocrine GnRH cells. Although *Gnrh1* expression is extinguished in these cells, *Rosa*-driven tdTomato expression persists through life, resulting in ectopic tdTomato-labeled cells, consistent with (Harris et al., 2014), who reported strong *Gnrh1*-Cre labeling within the lateral septum.

Taking advantage of this phenomenon, we were able to visualize ectopic tdTomato-labeled cells and tdTomato-labeled GnRH neurons, both PI4KA knockouts. Here, we compare tdTomato-expressing cells in the medial and lateral septum (Fig. 9, top panels) and in the OVLT region of the hypothalamus (Fig. 9, bottom panels) of infantile (*A* and *B*) and adult (*C* and *D*) control and knockout females, immunostained for GnRH. Red indicates tdTomato-positive cells only, and yellow indicates tdTomato+GnRH coexpressing cells. The lateral septum showed cells positive only for tdTomato, consistent with literature (Skynner et al., 1999), and these cells were not affected by PI4KA deletion as they remained comparable to control females from PND10 to adulthood in knockout females (Fig. 9, top panels). In contrast, the medial septum and OVLT region showed cells co-expressing tdTomato and GnRH, which regressed in infantile knockouts, and these areas were devoid of tdTomato cells in adult knockouts (Fig. 9, bottom panels). Together, this indicates that the morphological changes preceded the death of GnRH neurons and that PI4KA knockout is not universally deleterious, but rather specific for neuroendocrine GnRH cells.

**Figure 9.**
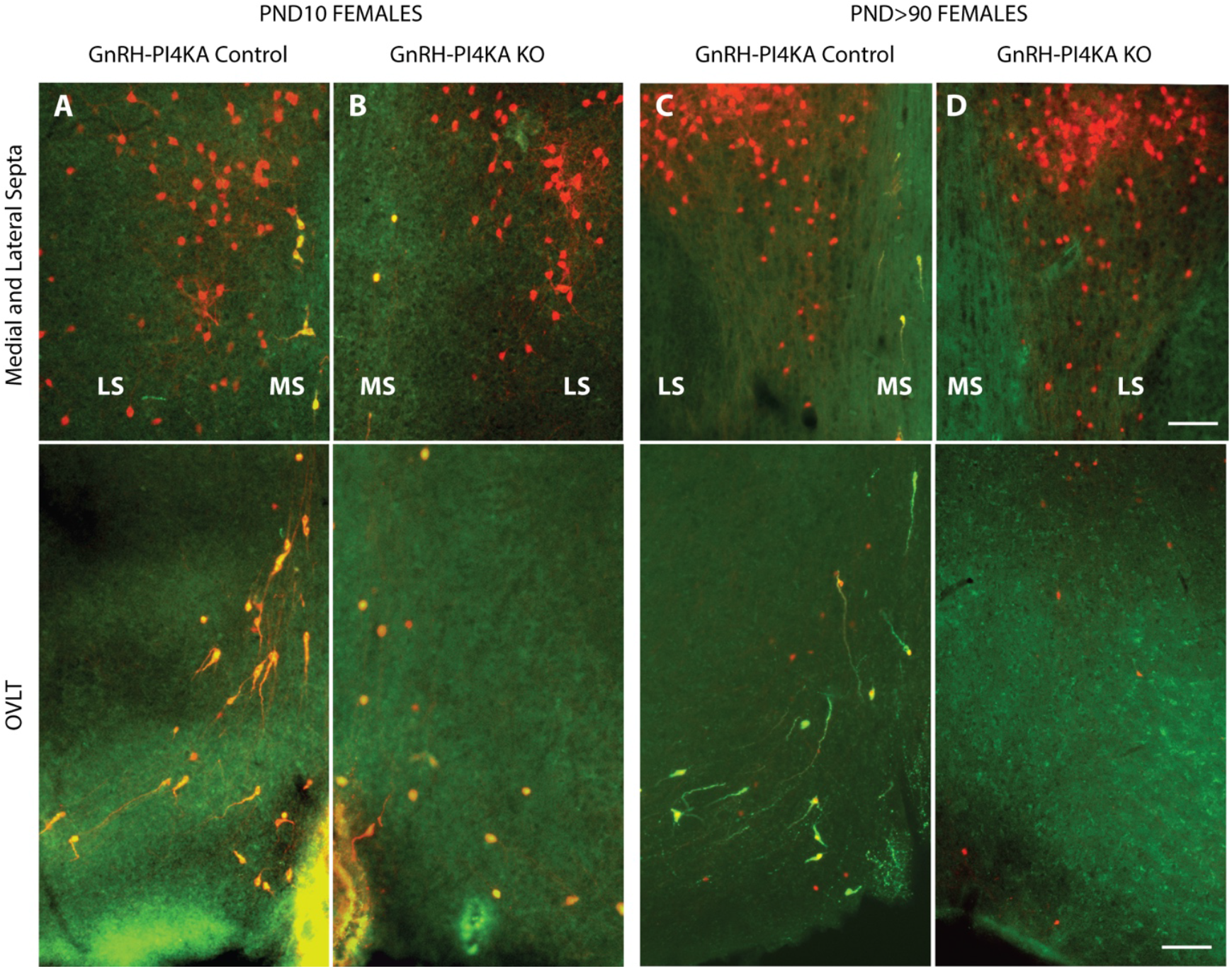
Comparison of tdTomato-expressing cells in the medial and lateral septa (upper panels) and in the OVLT region of the hypothalamus (lower panels) of infan0le (***A*** and ***B***) and adult (***C*** and ***D***) control and knockout female mice. Red indicates tdTomato-posi0ve cells, and yellow indicates tdTomato+GnRH-posi0ve cells. It should be noted that tdTomato-posi0ve neurons are present in the lateral septum (LS) and are no a?ected by PI4KA knockout (***A*** to ***D***, upper panels). tdTomato+GnRH-posi0ve cells present in the medial septa (MS) and OVLT region of the hypothalamus (***A*** and ***C***) show their bipolar morphology in controls, however these cells were round in infan0le knockouts (***B***), and disappeared in adult knockouts (***D***). Horizontal bars at 100 µm apply to all panels.

## Discussion

Here we show that PI4KA knockout in GnRH neurons causes infertility in both females and males, reflecting the absence of puberty and underdeveloped gonads and reproductive organs. Since GnRH controls the onset of puberty and fertility, these observations suggest a key role for PI4KA in the function of GnRH neurons. To further investigate this hypothesis, we performed immunolabeling studies, using adult controls and knockouts and a specific antibody against GnRH. These experiments revealed a lack of GnRH immunoreactivity in the OVLT area and the median eminence of the knockouts. We also observed a significant reduction in *Gnrh1* expression, as assessed by qRT-PCR of hypothalamic tissue in adult females and males.

Without GnRH, the expression of gonadotroph-specific genes should be reduced (Stamatiades and Kaiser, 2018). This in turn should reduce the synthesis and release of gonadotropins, leading to underdeveloped gonads, absence of puberty, and infertility (Coss, 2018). Accordingly, we observed a significant reduction in *Lhb* and *Gnrhr* expression in adult pituitaries from both female and male adult knockouts. Expression of *Pomc, Tshb*, and *Gh1*, a marker gene for corticotrophs/melanotrophs, thyrotrophs, and somatotrophs, respectively, was not significantly affected, further indicating the specificity of the knockout. Within pituitary cells, *Spp1* is expressed only in gonadotrophs (Fletcher et al., 2019), and expression of this gene was significantly reduced in knockouts. However, *Spp1* expression is not regulated by GnRH (Bjelobaba et al., 2019), but is stimulated by gonadal steroid hormones (Dunlap et al., 2008), which also stimulate *Prl* expression in lactotrophs (Grattan, 2015). It is therefore reasonable to conclude that the reduced expression of *Spp1* and *Prl* reflects the lack of gonadal steroidogenesis in knockout females and males.

Since GnRH secretion is regulated by kisspeptin (Kirilov et al., 2013; Han et al., 2015; Nandankar et al., 2021), it was important to compare the status of kisspeptin neurons in postnatal controls and knockouts. There are two major hypothalamic populations of kisspeptin neurons in rodents, one in the RP3V, and the other in the ARN (Goodman and Lehman, 2012). The RP3V kisspeptinergic population is sexually dimorphic, being larger in females than in males. In contrast, no sex differences were observed in the ARN kisspeptin neuron population (Kauffman et al., 2007). Here we show that the sexually dimorphic expression of RP3V kisspeptin is preserved in knockouts and that kisspeptinergic fibers were less abundant in same sex controls. This indicates that the sex steroid status of knockout females at PND15, although likely attenuated, is not as drastic as that of PND15-gonadectomy, which eliminates RP3V immunoreactivity (Clarkson et al., 2009). Furthermore, no sex-specific pattern of kisspeptin expression in ARN was observed in control and knockout groups. Knockout showed weak kisspeptin immunoreactivity of fibers with easily identified cell bodies, whereas controls showed strong kisspeptin labeling of fibers without visible cell bodies. In line with the literature (Smith et al., 2005a; Smith et al., 2005b), these data suggest that PI4KA knockout in GnRH neurons affects kisspeptin expression in a nucleus-specific manner due to the loss of gonadal steroidogenesis. This is consistent with the fact that GnRH is not required for the establishment of kisspeptin populations (Kim et al., 2013).

Previous studies have clarified that embryonic steps involve the development of GnRH neurons: from their differentiation in the olfactory placodes, their migration to the preoptic area and mediobasal hypothalamus, and their projection to the median eminence (Duittoz et al., 2022). The postnatal period is characterized by the establishment of the GnRH network with RP3V kisspeptin neurons for puberty onset (Clarkson et al., 2009), ARN kisspeptin neurons for the initiation of GnRH pulse generation in both females and males (McQuillan et al., 2019), and RP3V kisspeptin neurons for the preovulatory GnRH surge only in females (Clarkson et al., 2008).

Here we indirectly show that embryonic steps in the development of GnRH neurons are not critically affected by PI4KA knockout. Indeed, our qRT-PCR experiments with PND3 females and males revealed no difference in *Gnrh1* expression in the knockouts. Furhtremore, we show that distribution of GnRH neurons and their projections in the OVLT region and median eminence was normal in infatile knockout females and males. However, *Gnrh1* expression was significantly affected in knockout females and males by PND9. Furthermore, most GnRH cell bodies in PND10 mice lack processes, and immunoreactivity was somewhat reduced in the median eminence of knockouts, compared with age-matched controls. Such morphological changes, combined with a decrease in *Gnrh1* expression, indicate that loss of PI4KA begins to affect postnatal development and function of the GnRH system during the infantile period.

To better understand the postnatal changes in GnRH neurons of knockouts, we further characterized the developmental expression profile of *Gnrh1* in the rostral hypothalamus and the expression profiles of *Kiss1* in both the rostral and caudal hypothalamus of control and knockout groups. The main finding of these experiments was the progressive decline in *Gnrh1* expression with postnatal age in both females and males, as well as the absence of the postnatal region-specific changes in *Kiss1* expression in the rostral and caudal hypothalamus in knockouts, changes that are normally associated with the establishment of steroid feedback as mice progress through puberty. Furthermore, we show parallelism in *Tac2* and *Kiss1* expression in knockouts in the caudal hypothalamus containing ARN kisspeptin neurons.

Gonadal steroid hormones regulate GnRH secretion and GnRH neurons lack steroid receptors, suggesting their indirect effects. Further studies revealed that RP3V kisspeptin neurons are responsible for the positive feedback of estradiol, while ARN kisspeptin neurons mediate the negative feedback of estradiol and testosterone (Adachi et al., 2007; Yeo et al., 2014; Wang et al., 2018). Consistent with this, our data show that sex steroid feedback is not established as knockouts grow. *Kiss1* expression remains low in the rostral hypothalamus due to the lack of steroid stimulation and increases in the caudal hypothalamus due to the lack of steroid inhibition. It is noteworthy that *Tac2* expression follows the same profile as *Kiss1* expression, as the *Tac2* gene is regulated in the same way (Kauffman et al., 2009).

These experiments could not clarify the nature of *Gnrh1* loss: irreversibly silenced *Gnrh1* expression or caused by premature death of GnRH neurons. To study the fate of GnRH neurons postnatally, it was necessary to identify them independently of their peptidergic phenotype. This was done using tdTomato reporter reflecting *Gnrh1*-driven Cre recombination, as described in Material and Methods section. Once recombination occurs in target cells, tdTomato is present until their death regardless of the nature of postnatal differentiation, as it is driven by the constitutively active ROSA26 promoter (Srinivas et al., 2001). Colabeling of GnRH immunoreactive neurons with tdTomato confirmed the validity of our model for tracing GnRH neurons. The change in morphology of PND10 knockout GnRH neurons indicates the onset of degeneration of these cells. Furthermore, the loss of GnRH immunoreactivity in adult knockouts was accompanied by the loss of tdTomato labeling, indicating cell death.

In contrast, tdTomato cells in the lateral septum, which reflect transient expression of *Gnrh1* during development (Skynner et al., 1999), i.e. ectopic tdTomato cells, persisted throughout life and were apparently unaffected by PI4KA knockout. As in GnRH neurons, PI4KA knockout in mouse gonadotrophs also caused infertility. However, in this case, infertility was not due to cell death, but to impaired GnRH receptor signaling, which requires PI(4,5)P2 production (Constantin et al., 2023). Impaired GnRH receptor signaling led to dedifferentiation of existing gonadotrophs and reduced postnatal differentiation of new gonadotrophs from marginal zone stem cells (Smiljanic et al., 2025). Therefore, cell death by PI4KA knockout is novel finding for neuroendocrine cells.

The relationship between phosphoinositides and cell death is a very complex issue and encomnpasses both the action of phosphoinositides as second messengers and their roles in the actin cytoskeleton and membrane dynamics, as well as their direct role in modulating cell death pathway (Phan et al., 2019). For example, phosphatidylinositol-3-kinase has been suggested to inhibit apoptosis (Webb et al., 2000), but to accelerate necrotic cell death (Aki et al., 2001). More specifically, GnRH neuron survival has been shown to depend on semaphorin 3E signaling, i.e. activation of phosphatidylinositol-3-kinase pathway (Cariboni et al., 2015). Lipotoxic disruption of NHE1 interaction with PI(4,5)P2 also accelerate apoptosis of proximal tubules (Khan et al., 2014), as does knockdown of PI4K2A or PI4KB (Chu et al., 2010). Dysregulation of PI4P production by knockout of PI4KA is also associated with increased lysosome-dependent cell death (Xu et al., 2025). Thus, the loss of GnRH neurons in prepubertal animals due to PI4KA knockout is consistent with preliminary findings in other cell types, suggesting a direct or indirect cell type-specific role of PI4KA-derived phosphoinositides in the postnatal survival and function of these neuroendocrine cells.

## Acknowledgments

We are thankful to Prof. Greg Anderson for the GA02 guinea pig anti-GnRH antibody. The floxed PI4KA mouse strain was provided by GlaxoSmithKline LLC under a Materials and Cooperative Research and Development Agreement. This research was supported by the Intramural Research Program of the National Institutes of Health (NIH). The contributions of the NIH author(s) are considered Works of the United States Government. The findings and conclusions presented in this paper are those of the authors and do not necessarily reflect the views of the NIH or the U.S. Department of Health and Human Services.

